# Heterochronically expressed midline netrin was recruited to guide mesoderm migration in epibolic gastrulation of the leech

**DOI:** 10.1101/803353

**Authors:** Jun-Ru Lee, Dian-Han Kuo

## Abstract

Netrin is a remarkably conserved midline landmark, serving as a chemotactic factor that organizes the bilateral neural architecture in the post-gastrula bilaterian embryos. Netrin signal also guides cell migration in many other neural and non-neural organogenesis events in later developmental stages, but it has never been before found to participate in gastrulation – the earliest cell migration in metazoan embryogenesis. Here, we found that netrin signaling molecules and their receptors are expressed during gastrulation of the leech *Helobdella*. Intriguigly, *Hau-netrin-1* was expressed in the N lineage, the precursor of ventral ectoderm, at the onset of gastrulation. We demonstrated that the N lineage is required for the entrance of mesoderm into the germinal band and that misexpression of Hau-netrin-1 in early gastrulation prevented mesoderm from entering the germinal band. Together, these results suggested that Hau-netrin-1 secreted by the N lineage guides mesoderm migration during germinal band assembly. Furthermore, ectopic expression of Hau-netrin-1 after the completion of germinal band assembly disrupted the epibolic migration of the germinal bands in a later stage of gastrulation. Thus, Hau-netrin-1 is likely involved in two distinct events in sequential stages of leech gastrulation: the assembly of germinal bands in early gastrulation and their epibolic migration in mid-gastrulation. This mode of gastrulation as observed in the leech is apomorphic for clitellate annelids. We postulated that a heterochronic shift of netrin gene expression in the clitellate ancestor might have facilitated the evolutionary emergence of a novel form of gastrulation in this lineage.

## Introduction

Gastrulation is the earliest and most fundamental morphogenetic process in the development of the animal body plan. Nevertheless, the patterns of cell migration in gastrulation can vary greatly among species sharing the same body plan, and such variation has often been ultimately attributed to the life history strategies adopted by individual species (Collazo et al., 1994; del Pino and Elinson, 1983; Wray and Raff, 1991). Beyond the ecological drives, however, molecular changes underpinning the evolutionary diversification of gastrulation remain elusive. Among the annelids, a unique form of gastrulation, in which posterior growth and axial patterning proceed simultaneously during epibolic migration of the primordial trunk tissue, has evolved in the clitellate lineage (earthworms and leeches) (Kuo, 2017). Clitellates are an ecologically successful group in the freshwater and terrestrial habitats, and the unique mode of embryonic development is likely among the key innovations that enabled the clitellate ancestor to invade the unoccupied land habitats and undergo adaptive radiations (Kuo, 2017). By comparing embryology of the clitellates and the marine polychaete annelids, direct development was proposed to arise by the acceleration of trunk development in clitellates (Kuo, 2017).

Netrin is a chemotactic factor involved in orchestrating tissue morphogenesis in a great variety of organogenesis events (Hinck, 2004; Sun et al., 2011; Ziel and Sherwood, 2010). Among its many functions in animal development, netrin is best known as an axon guidance molecule marking the midline of the central nervous system (CNS) in flies and vertebrates (Harris et al., 1996; Kennedy et al., 1994; Mitchell et al., 1996; Serafini et al., 1996), and the expression of netrin along or near the CNS midline is largely conserved across Bilateria (Denes et al., 2007; Harris et al., 1996; Hotta et al., 2000; Kugler et al., 2011; Linne and Stollewerk, 2011; Lowe et al., 2006; Mitchell et al., 1996; Shimeld, 2000; Wadsworth et al., 1996). Interestingly, despite the absence of a bilaterally symmetric CNS, netrin also exhibited a polarized expression along the directive axis in the cnidarian planula larva (Matus et al., 2006), implying that netrin may be an ancient component in the patterning system for the second body axis. Therefore, netrin is a fitting candidate for testing our hypothesis of accelerated trunk development in the clitellates.

The earliest site of netrin expression is the CNS midline in the species that have been examined thus far, and therefore netrin has never been implicated in gastrulation – the earliest cell migration event in an animal embryo. Here, we unveil an unprecedented case for netrin-guided cell migration during gastrulation of the leech *Helobdella*. At the onset of gastrulation, *Hau-netrin-1*, one of the two netrin homologs in the leech, was expressed in the N lineage, among whose progeny is the ectodermal midline. In a later stage of gastrulation, *Hau-netrin-2* was also turned on in cells that would become the mesodermal midline after the completion of gastrulation. These findings indicated a possible heterochronic shift in the developmental expression of both leech netrin genes relative to the bilaterian archetype. Furthermore, we demonstrated that the early expression of Hau-netrin-1 in the N lineage is involved in two distinct cell migration events in leech gastrulation: first, the assembling of mesoderm into the germinal band in early gastrulation, and second, the epibolic migration of the germinal band in mid-gastrulation; both are unique features in clitellate gastrulation. Together, our results suggested that the heterochronically expressed netrin was recruited to regulate the formation and migration of germinal band, which is central to the evolution of direct development and a new mode of gastrulation in the clitellate lineages.

## Results

### Gastrulation of the leech *Helobdella*

The pattern of cell movement during gastrulation is generally conserved among the clitellate embryos (Anderson, 1973). Briefly, the left and right germinal bands, giving rise to the trunk ectoderm and mesoderm, elongate by posterior addition from the teloblast stem cells and migrate over the circumference of the embryo to enclose the endoderm. Not only the morphogenetic process, the patterning of cell fates in the clitellate gastrula, which is dominated by a stereotyped cell lineage – leading from individual teloblasts to the differentiated ectodermal and mesodermal components of the segmented trunk, is also evolutionarily conserved (Goto et al., 1999; Storey, 1989; Weisblat and Shankland, 1985). These traits are clitellate apomorphies and critical for the evolution of direct development in clitellate annelids (Kuo, 2017).

Among the clitellates, the cellular and molecular basis of embryonic development are best characterized in the leech *Helobdella* (Weisblat and Kuo, 2009). Developmental events leading to the completion of gastrulation are summarized below. After fertilization, the mRNA-enriched teloplasm emerged in the zygote and then segregated into the D quadrant in the first two cleavages (stages 1-3). During stages 4-6, series of spiral and bilateral cleavages generate dozens of micromeres near the animal-pole region, three large endoderm-fated macromeres on the vegetal side of the embryo, and five bilateral pairs of teloblast stem cells; each cell can be identified by its unique size and position (Weisblat and Huang, 2001; Weisblat and Kuo, 2014). Among these, teloblasts inherit the teloplasm from the D quadrant and give rise to the segmented trunk ectoderm and mesoderm (Figure 1A).

**Figure 1.**
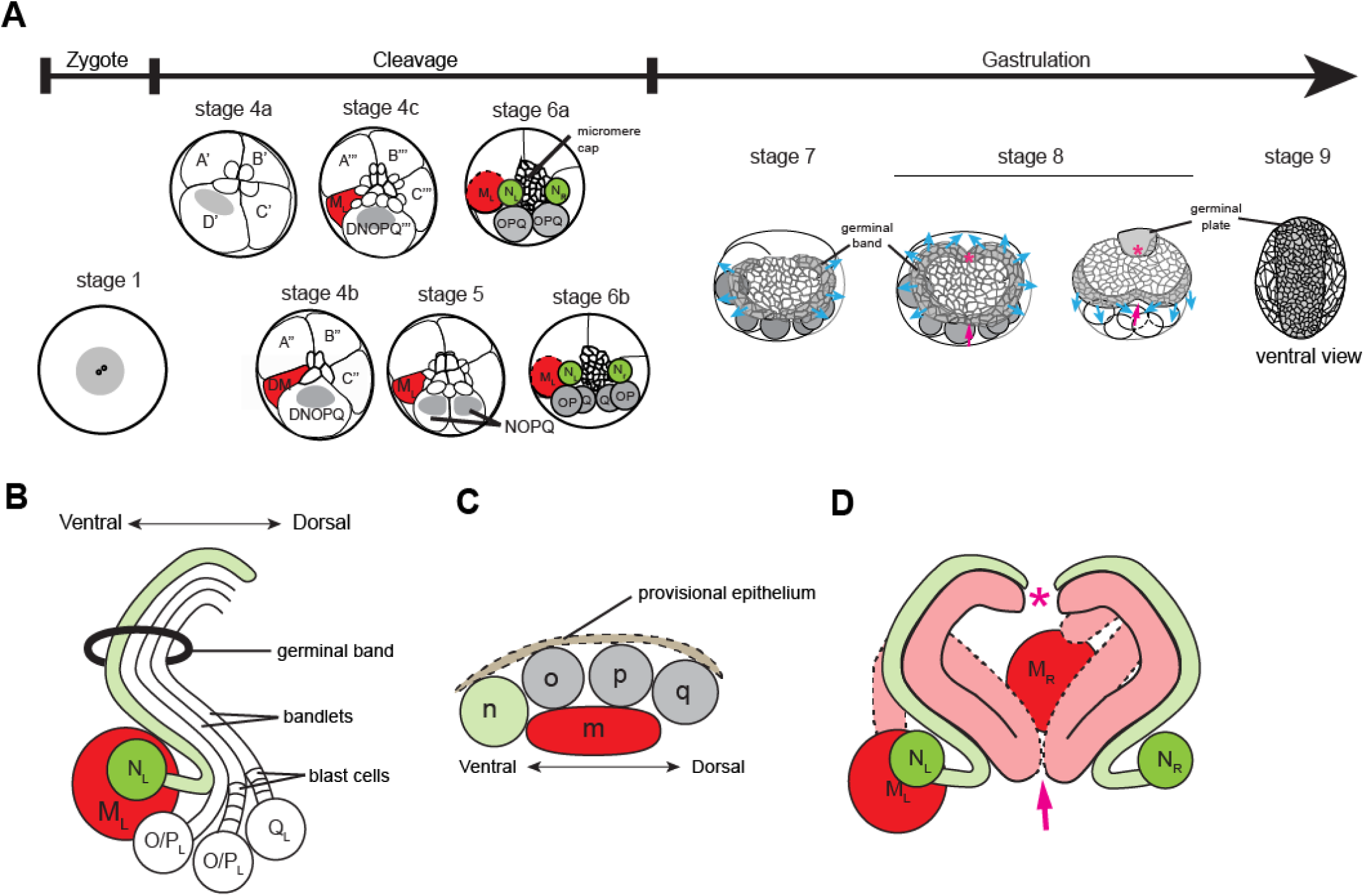
Schematic depictions of germinal band development during leech gastrulation. (A) Early developmental events leading to the completion of gastrulation in the leech embryo. Spiral cleavages take place in stage 4; the teloblastogenic bilateral cleavages are in stages 5 and 6; gastrulation and axial elongation by the posterior growth of germinal band tissue are in stages 7 and 8. The left and right germinal bands emerge during stage 7; the germinal bands undergo epibolic migration to form the germinal plate on the ventral surface during stage 8. Blue arrows indicate the directions of germinal band migration in stage 8. The completion of gastrulation defines the onset of stage 9. All drawings, except for that of stage-9 embryo, is in animal-pole/dorsal view. The stage-9 embryo is in the ventral view. (B) Germinal band tissue arises from the asymmetric teloblast divisions (see text for details). In a bandlet, older blast cells are anterior to younger blast cells of the same teloblast lineage, and the youngest blast cells are in the proximity of the teloblasts. (C) Cross-section of a leech germinal band. The m bandlet lies underneath the four dorsoventrally arranged ectodermal bandlets, and a thin layer of provisional epithelium covers the germinal band. (D) The organization of the left and right m bandlets relative to the ectodermal n bandlets in early stage 8. The other ectodermal bandlets (o, p, and q) are removed for clarity. The posterior portion of the m bandlet folds in a stereotyped pattern in the narrow coelomic space. The mesoderm lineage is labeled in red; the N lineage, which gives rise to the ventralmost portion of trunk ectoderm, is in green; the remaining ectodermal teloblasts and teloplasm in early blastomeres are in grey. Pink asterisks mark the anterior end of the germinal band/plate; pink arrows mark the posterior end of the germinal band, where the individual bandlets converge to form the germinal band.

Of the five teloblast lineages, the mesodermal M lineage arises from the DM blastomere, the vegetalward daughter of the D’ macromere, whereas the four ectodermal lineages (N, O, P, and Q) arises from the DNOPQ blastomere, the animalward daughter of D’ (Figure 1A). Each teloblast undergoes iterated asymmetric cell divisions to produce a ‘bandlet’ of segment founder cells, or ‘blast cells’ (Figure 1B); bandlet and blast cells are denoted with the lower case letter representing the teloblast lineage from which they originate. The five ipsilateral bandlets converge at the posterior end of the embryo and are assembled into the germinal bands starting at stage 7 (Figure 1A). In the germinal band, the mesodermal bandlet lays underneath the four ectodermal bandlets, with n, o, p, and q, arranged from ventral to dorsal (Figures 1C and 1D). Blast cells in a given lineage then follow a specific program of cell divisions to give rise to a definitive set of differentiated pattern elements in stage 9 and beyond (Weisblat and Shankland, 1985; Zackson, 1984). During stage 8, the elongating germinal band undergoes a ventralward epibolic migration across the circumference of the embryo as the newly born blast cells join the germinal band from the posterior, and the micromere cap between the two germinal bands also expands by cell proliferation (Figure 1A). Eventually, the left and right germinal bands meet at the prospective ventral midline to form the germinal plate (Figure 1A). The expansion of the micromere cap and the epibolic movement of germinal bands in stages 7 and 8 constitute the gastrulation of the leech (Smith et al., 1996; Smith and Weisblat, 1994), and the completion of the coalescence of the left and right germinal band along the entire body length marks the end of gastrulation and the onset of stage 9.

### *Netrin* homologs are expressed in specific cell populations beginning at the onset of leech gastrulation

The expression pattern of a netrin gene in the developing CNS was previously characterized in the medicinal leech *Hirudo* and is in agreement with the conventional idea that it serves as the midline landmark that guides the morphogenesis of CNS (Aisemberg et al., 2001; Gan et al., 1999). Nonetheless, netrin expression in earlier developmental stages remained unknown for the leech. Given that the movement of germinal band tissue in gastrulation is highly coordinated and stereotyped, it is possible that guiding molecules may participate in some of these events. To explore such a possibility, we characterized the expression patterns of netrin genes during gastrulation of the leech *Helobdella*, whose early embryos are more experimentally tractable than that of *Hirudo*. We identified two netrin homologs in the whole genome assembly of *Helobdella robusta* (Table S1) and cloned *Hau*-*netrin-1* and *Hau-netrin-2* from the sibling *H. austinensis*. *Hau-netrin-1* is the ortholog of previously identified *Hirudo netrin*; *Hau-netrin-2* is more derived in levels of both molecular sequence and domain structure. Based on the phylogenetic analysis of the conserved laminin-like domain, the two leech netrin genes were likely resulting from a gene duplication event in the ancestor of clitellates or leeches (Figure S1A).

We next probed their expression patterns in *Helobdella* embryo during gastrulation (stages 7 and 8) by using whole-mount *in situ* hybridization (WMISH). Expression of *Hau*-*netrin-1* was restricted to the n bandlets and the opq’ micromeres in early stage 7 (Figures 2A and 2B). In late stage 7, *Hau-netrin-1* expression disappeared in the micromeres but persisted in the n bandlet (Figures 2C and 2D). In stage 8, as the n blast cell clone continues to divide, *Hau-netrin-1* expression becomes further restricted to a specific set of N sublineages (Figures 2E and 2F). Cells expressing the highest level for *Hau-netrin-1* converged at the ectodermal midline (Figure 2E), and they are likely precursors of the neuropile gila that constitute the midline of the ventral nerve cord (Bissen and Weisblat, 1987). In stage 9 and beyond, the expression pattern of *Hau-netrin-1* (Figures S2A-S2D) was comparable to that of *Hirudo* netrin (Aisemberg et al., 2001; Gan et al., 1999), consistent with their orthologous relationship.

**Figure 2.**
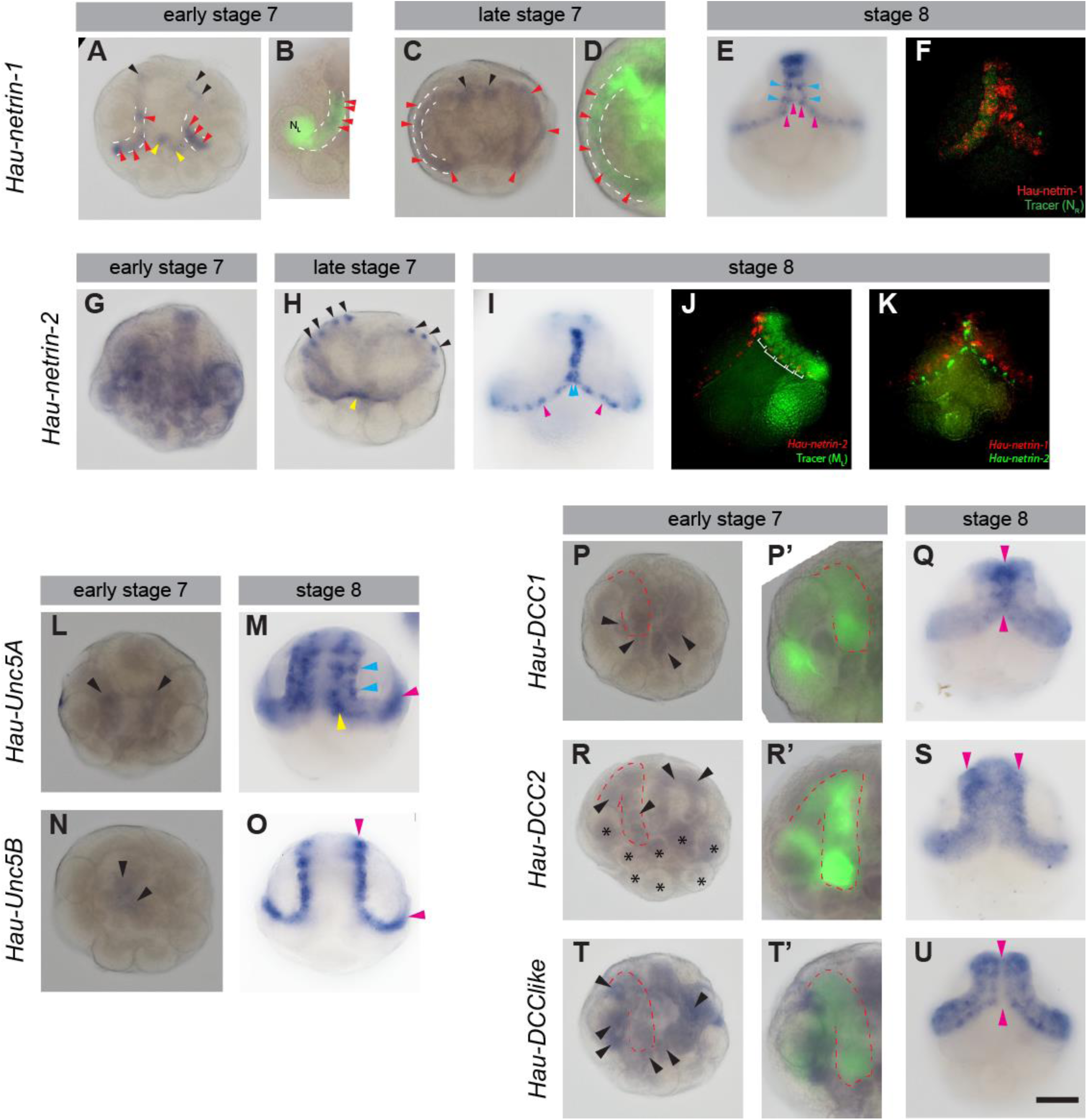
Expression patterns of the netrin ligands and receptors during leech gastrulation. (A) In early stage 7, *Hau-netrin-1* is expressed in the n bandlets, a pair of posterior micromeres (yellow arrowheads), and a few cells at the anterior margin of the micromere cap (black arrowheads). Red arrowheads indicate the N-lineage cell expressing *Hau-netrin-1*, and white dashed lines outline the n bandlets in panels A-D. (B) WMISH of *Hau-netrin-1* in a stage-7 embryo, whose N_L_ teloblast was injected with a tracer in stage 6a. (C) In late stage 7, *Hau*-*netrin-1* expression (red arrows) remained in the n bandlets (white dashed lines) and bilateral clusters of anterior micromeres (black arrows). (D) Tracer labeling identified the n bandlets (white dashed lines) with positive *Hau-netrin-1* WMISH signal (red arrows). (E) In stage 8, *Hau-netrin-1* was up-regulated in specific cells located in the ventral region of the n blast cell clones (pink arrowheads); *Hau-netrin-1* was up-regulated in additional cells in older n clones within the germinal plate (blue arrowheads). (F) Fluorescent *Hau-netrin-1* WMISH (red) using a tracer labeled embryo confirmed its restricted expression (red) in the N lineage (green). (G) WMISH revealed that *Hau-netrin-2* is broadly expressed in early stage 7. (H) *Hau-netrin-2* expression is restricted to a few cells near the anterior margin of the micromere cap on the left and right sides (black arrowheads) and at the posterior border of the micromere cap (yellow arrowhead) in late stage 7. (I) *Hau-netrin-2* expression shifts to the prospective mesoderm midline cells (pink arrowheads) in the germinal band and become highly expressed along the mesodermal midline (blue arrowheads) of the germinal plate in stage 8. (J) Fluorescent *Hau*-*netrin-2* WMISH (red) confirmed its expression within the mesoderm of a stage-8 embryo, whose M_L_ teloblast was injected with tracer (green) in stage 6. (K) Double fluorescence WMISH of *Hau-netrin-1* (red) and *Hau-netrin-2* (green) in a stage-8 embryo. (L) WMISH of *Hau-Unc5A* revealed weak *Hau-Unc5A* expression in the left and right flanks of the micromere cap (black arrowheads) in early stage 7. (M) WMISH revealed *Hau-Unc5A* expression in the M lineage within the germinal band (pink arrowhead) and in clusters of ectodermal cells in the dorsal (blue arrowheads) and lateral (yellow arrowhead) regions of the germinal plate in stage 8. (N) WMISH revealed weak *Hau-Unc5B* expression in the center of the micromere cap (black arrowheads) in early stage 7. (O) *Hau-Unc5B* is expressed in the dorsal edge of the germinal bands and germinal plate (pink arrowheads) in stage 8. (P) WMISH revealed that *Hau-DCC1* is expressed in selected superficial cells (black arrowheads) in the posterior region of the micromere cap in stage 7. In the panel and panels R and T, the red dashed line outlines the left m bandlet, which is visualized by tracer labeling shown in panels P’, R’, and T’, respectively. (Q) *Hau-DCC1* was upregulated in the ventral region of the germinal plate (pink arrowheads) in stage 8. (R) *Hau-DCC2* was expressed in the m bandlet (black arrowheads) as well as in other teloblast lineages (black asterisks) in stage 7. (S) WMISH revealed *Hau-DCC2* up-regulation in the dorsal region of the germinal plate (pink arrowheads). (T) *Hau-DCClike* was expressed in the peripheral region of the micromere cap (black arrowheads) in stage 7. (U) *Hau-DCClike* was expressed in ectodermal bandlets, except for the midline region of the germinal plate (pink arrowheads), in stage 8. WMISH was performed in combination with tracer labeling to identify the lineage identities of the cells that express a specific gene. All stage-7 embryos are shown in animal-pole/dorsal view; all stage-8 embryos are shown in the ventral view; anterior is to the top in all panels. Scale bar: 150 μm in panels A, C, E-U; 100 μm in panels B, D, P’, R’, and T’.

The expression of *Hau-netrin-2* was highly dynamic during development. In early stage 7, *Hau-netrin-2* was broadly expressed in the micromere cap, teloblasts, and blast cells (Figure 2G). Near the end of stage 7, however, *Hau-netrin-2* expression resolved into a few cells at the edge of the micromere cap when epibolic migration of the germinal bands began (Figure 2H). The sites of *Hau-netrin-2* expression changed yet again in the middle of stage 8, as the *Hau*-*netrin-2* transcript was detected in the ventralmost mesodermal sublineages in the migrating germinal bands (Figures 2I and 2J). These *Hau-netrin-2*-expressing cells became the mesodermal midline after the left and right germinal bands converge to form the germinal plate. Double fluorescent WMISH showed that cells expressing *Hau-netrin-2* in the mesoderm migrated ahead of the ectodermal *Hau-netrin-1*-expressing cells in the germinal bands (Figure 2K). In stage 9, *Hau-netirn2* expression continued in the ventral midline of the body-wall mesoderm and visceral mesoderm and was turned on in the developing nephridia (Figures S2E-S2H). In stage 11, *Hau-netrin1* and *Hau-netrin-2* were expressed in complex patterns (Figures S2D and S2I), suggestive for their combinatorial roles in organogenesis of the proboscis, intestinal caecum, and ventral midline structures.

### Netrin receptors are expressed during gastrulation

Netrin signals through distinct types of receptors to elicit cellular response (Cirulli and Yebra, 2007). To determine whether the netrin signal operates during leech gastrulation, we next set out to examine the expression patterns of netrin receptors in stage-7 and -8 embryos. Two UNC5-type and three DCC-type receptors were identified in the *Helobdella* genome (Figures S1B and S1C; Table S1). Distinct expression patterns were detected for *Hau-Uuc5A* and *Hau-Unc5B* in the micromere cap during stage 7 (Figures 2L and 2N). In stages 8, *Hau*-*Unc5A* was expressed in ectodermal cell clusters within the ventrolateral and dorsal regions of the converging germinal bands (Figure 2M), whereas *Hau-Unc5B* was expressed at the dorsal edge of the mesodermal bandlet (Figure 2O). Among the three DCC-type receptors, *Hau*-*DCC1* expression was detected in a few epithelial cells of the micromere cap during stage 7 (Figures 2P) and then in the germinal bands and the germinal plate during stage 8, with the highest expression level detected in the ventral region (Figure 2Q). *Hau-DCC2* expression was restricted to the m bandlet in stage 7 (Figure 2R), and it became highly expressed in the dorsal territory of the germinal plate (Figure 2S), complementing the expression pattern of *Hau*-*DCC1*. The expression of *Hau-DCClike* was observed in the peripheral of the micromere cap in stage 7 (Figure 2T) and became broadly distributed in the ectodermal bandlets in stage 8 (Figure 2U). Tracer labeling confirmed the expression of *Hau-DCC2* in the m bandlet (Figure 2R’) and showed that *Hau-DCC1* and *Hau-DCClike* were not expressed in mesoderm during stage 7 (Figures 2P’ and 2T’). The expression pattern of each receptor in stage 9 is similar to the respective stage-8 pattern (Figures S2J-S2N).

To summarize, the expression of netrin receptors begins as early as in stage 7, and their expression became fairly specific and robust in stage 8, implying that netrin signaling can operate during leech gastrulation. Given the complexity in the dynamic expression of the netrin ligands and receptors, it is not possible to characterize all aspects of netrin’s involvement in the morphogenesis of leech embryo in this study. It was previously shown that the m bandlets undergo dramatic active migration in the assembling of germinal bands during stage 7 (Gline et al., 2011), and the specific expression of *Hau-DCC2* in the m bandlet raised the possibility that the netrin signal is involved in the migration of the m bandlets. Therefore, we first focused on mesoderm migration.

### The n bandlet guides the entrance of mesodermal bandlet into the germinal band

Previous blastomere ablation experiments revealed that interaction with ectoderm is required for proper positioning of the m bandlets in the early stages of germinal band assembly (Blair, 1982). Since *Hau-netrin-1* was expressed in the n bandlet, we hypothesized that Hau-netrin-1 from the n bandlet might mediate this ectoderm-mesoderm interaction that guides the mesoderm migration. In that case, the deletion of the n bandlet was expected to yield a similar result to the previous experiments in which the complete set of ectodermal bandlets was ablated. To test this hypothesis, we injected DNase together with a tracer into the N_L_ teloblast immediately after its birth to prevent further cell division in the N_L_ lineage and then injected a different tracer into the ipsilateral M_L_ teloblast for visualizing mesoderm migration. In 17% of the operated embryos, the labeled m bandlet entered the contralateral germinal band (Figures 3A and 3D), whereas we never observed this phenotype in the control embryos injected with tracer only in the N_L_ teloblast (Figures 3B and 3D). Note that ablation had not induced ectopic expression of *Hau-netrin-1* in any other lineage (Figure 3C). These results suggested that the n bandlet indeed contributed to the correct entrance of the m bandlet into the germinal band. However, the low penetrance rate suggested that additional factors might also help to guide the m bandlet into the germinal band.

**Figure 3.**
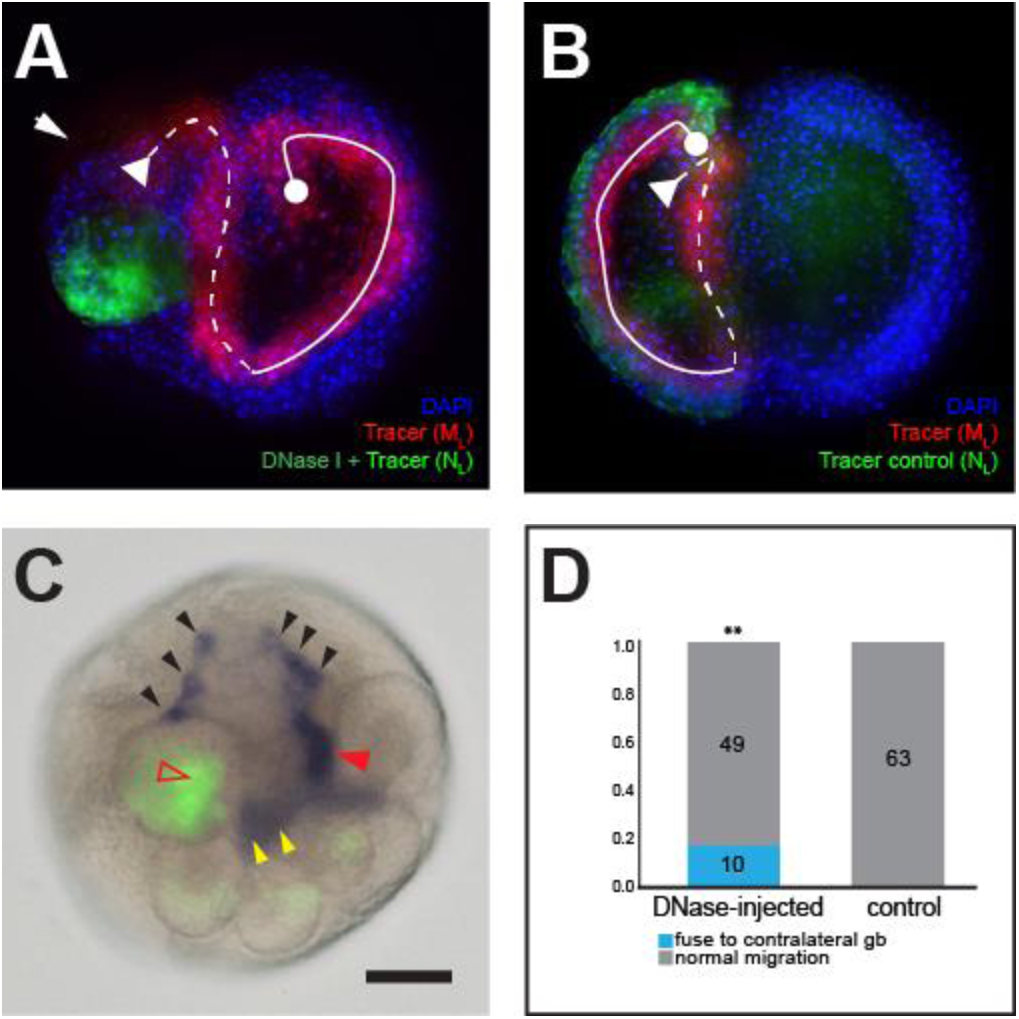
The N lineage is required for the normal positioning of the ipsilateral m bandlet in the germinal band. (A) The bandlet arising from the M_L_ teloblast (red) enters the contralateral, right germinal band after the ablation of left N lineage (green). The N_L_ teloblast was then injected with a mixture of fluorescein dextran and DNase immediately after its birth in stage 6a, and the M_L_ teloblast was next injected with a rhodamine dextran tracer. (B) The normal position of the left m bandlet (red) in a control embryo. (C) *Hau-netrin-1* expression was lost in the left n bandlet after its ablation (red open arrowhead); *Hau-netrin-1* expression in the right n bandlet (red arrowhead) appears normal. *Hau-netrin-1* expression in the posterior micromeres (yellow arrowheads) and anterior micromeres (black arrowheads) was bilaterally symmetric. (D) Quantification of the experiments shown in panels A and B. This switch-side phenotype was observed in 17% of the 59 operated embryos but was never observed in control embryos. Despite a low penetrance, the effect of N ablation on the position of the ipsilateral m bandlet was statistically significant. In panels A and B, the m bandlet is labeled with a white line; the dashed line represents the posterior region of the bandlet, which is yet to enter the germinal band; the solid-line portion represents the anterior region of the bandlet, which has already entered the germinal band; the dot at one end of the line marks the anterior tip of the bandlet; the triangle at the other end marks the posterior extreme of the bandlet, where the blast cells arise from the teloblast. Scale bar: 100 μm in panels A-C.

To document the normal interaction between the n and m bandlets during germinal band assembly, we went on to examine the developmental time course of double-labeled embryos whose N_L_ and M_L_ teloblasts were injected with different tracers. It was previously reported that the stage-7 embryo, the n bandlet is superficially localized, while the m bandlet runs across the interior of the embryo along the blastocoel wall during stage 7 (Gline et al., 2011). In early stage 7 (46 hrs AZD), the m bandlet, coming toward the anterior end of n bandlet from the interior, is perpendicular to the n bandlet, and their anterior tips contact each other under the micromere cap (Figures 4A). Eight hours later (54 hours AZD), the m bandlet moved toward the n bandlet progressively along the prospective anteroposterior axis, resulting in a hairpin bending of the m bandlet at the posterior edge of the micromere cap (Figures 4B). By 64 hours AZD, as the germinal bands elongate, the n and m bandlets migrate ventrally together in a coherent germinal band (Figures 4C). Therefore, the m bandlet is assembled into the germinal band by a zipping movement toward the superficial n bandlet (Figure 4D).

**Figure 4.**
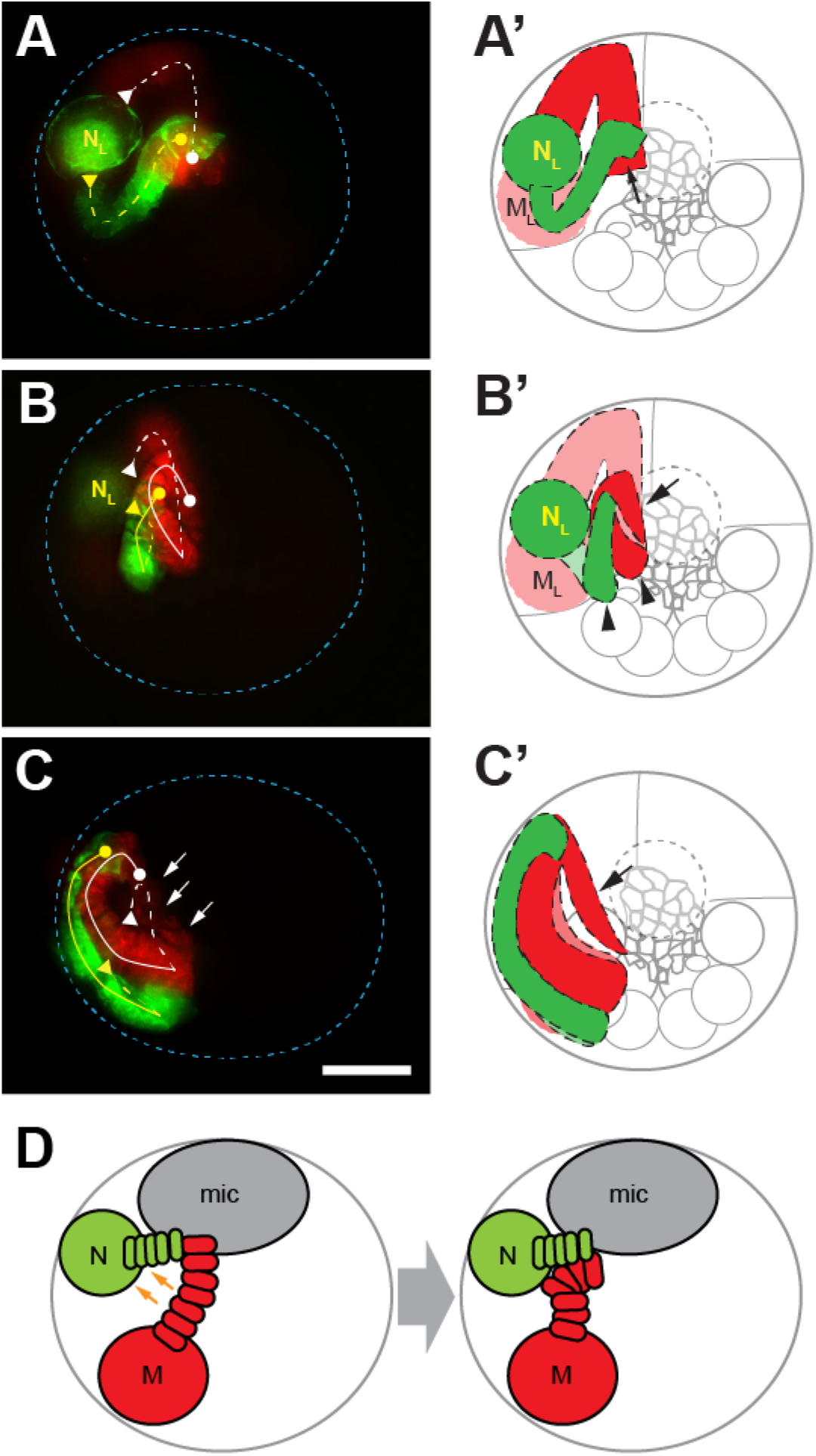
Time series analysis reveals morphogenetic movements of the m and n bandlets during germinal band assembly. (A) In early stage 7 (46 hours after zygote deposition; AZD), the N and M lineages contact each other at the anterior region of the bandlets. The image trace of the embryo in panel A is shown in panel A’. (B) At 54 hours AZD, the anterior cells of the n and m bandlets converge to from the germinal band. Both bandlets make a sharp U-turn at the posterior end of the bandlet. Trace of the embryo in panel B is shown in panel B’. (C) In late stage 7 (62 hours AZD), more cells in the n and m bandlets came into the germinal band as the germinal band migrates ventrovegetally. Arrows point to the migrating freckle cells derived from the anteriormost em_1_ and em_2_ blast cells of the M lineage. Trace of the embryo image in panel C is shown in panel C’. (D) Schematic representation of the process by which the m bandlet is assembled into the germinal band (see text for details).The left m bandlet is labeled with a white line and the left n bandlet with a yellow line; the dashed line represents the posterior region of the bandlet, which is yet to enter the germinal band; the solid-line portion represents the anterior region of the bandlet, which has already entered the germinal band; the dot at one end of the line marks the anterior tip of the bandlet; the triangle at the other end marks the posterior extreme of the bandlet, where the blast cells arise from the teloblast. The embryos were injected with rhodamine dextran tracer in the M_L_ teloblast right after the birth of the M teloblasts in early stage 5 and then with fluorescein dextran in the N_L_ teloblast in early stage 6a. The blue dashed lines mark the outlines of embryos. Scale bar: 150 μm.

### Hau-netrin-1 guides mesoderm migration during germinal band assembly

To see if Hau-netrin-1 is sufficient to guide the movement of the m bandlet, we next misexpressed Hau-netrin-1 in the m bandlet by co-injecting mRNA and a fluorescent tracer into an M teloblast immediately after its birth. This operation should flood the m bandlet with oversaturated netrin signal and therefore disrupt its oriented migration if Hau-netrin-1 can indeed guide mesoderm migration. As expected, in 85% of operated embryos, the m bandlet was detached from the germinal band and became “wandering” in the interior of the embryo (Figures 5A and 5D), whereas the m bandlet made an entrance into the germinal band normally in all control embryos (Figures 5B and 5D). By labeling the ectodermal bandlets with a different tracer, we were able to show that the mispositioned m bandlet failed to join the ectodermal bandlets (Figures 5E and 5E’). In contrast, the m bandlet was overlaid by the ectodermal bandlets along the entire length of the germinal band in all control embryos (Figures 5F and 5F’). These results suggested that the “wandering mesoderm” phenotype was caused by a failure in the entrance of the mesoderm into the germinal band during its assembly. The morphology of the m bandlet and the germinal band in embryos injected with *Hau-netrin-2* mRNA in the M teloblast appeared normal (Figures 5C and 5D), indicating that Hau-netrin-1, not Hau-netrin-2, can affect the migration of the m bandlet.

**Figure 5.**
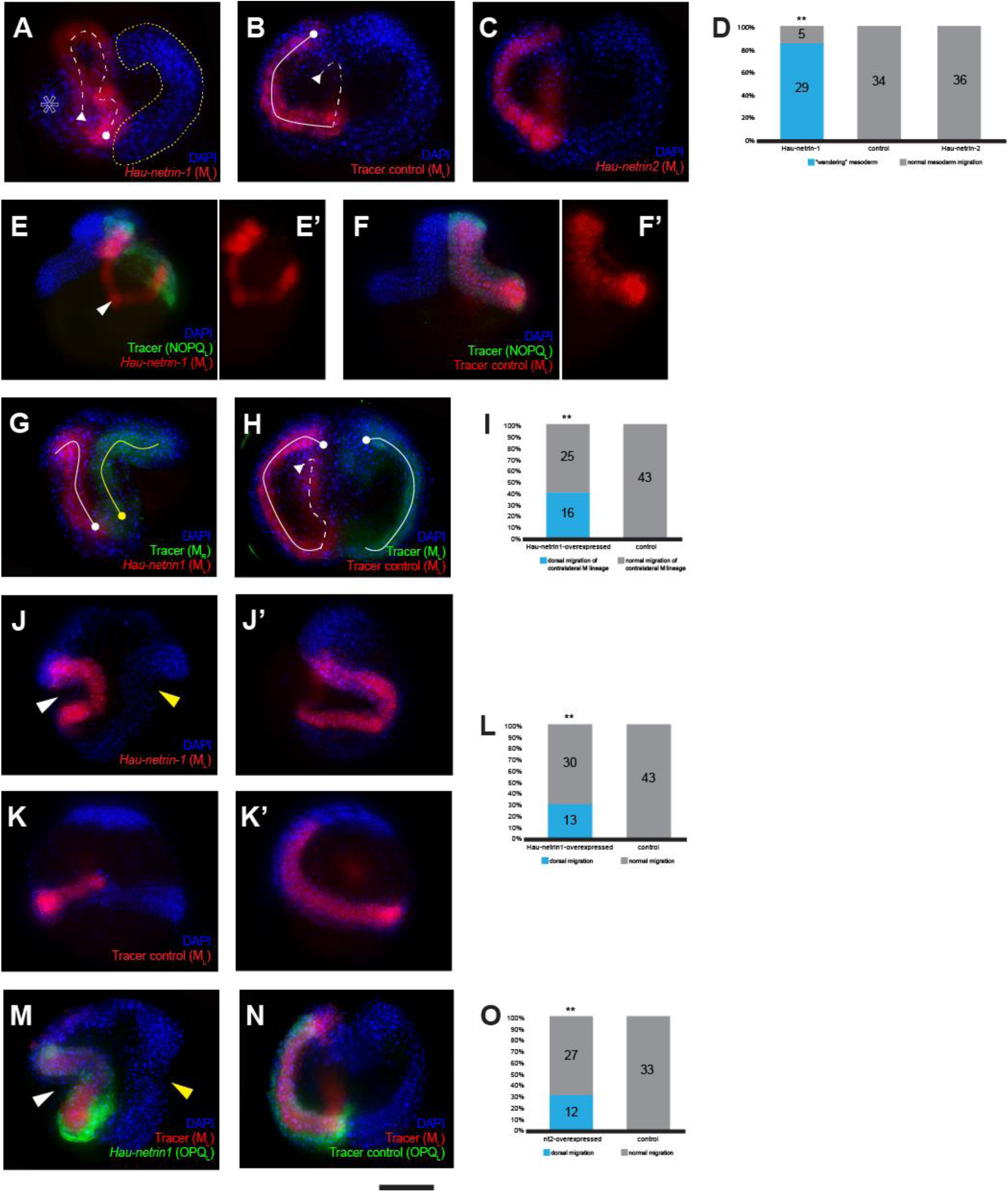
Mis-expression of Hau-netrin-1 prevented the m bandlet from entering the germinal band and disrupted the epibolic migration of the germinal band. (A) The left M lineage (red; labeled with white dashed line), which misexpresses Hau-netrin-1, was wandering in the coelomic space and failed to enter the germinal band. The anterior tip of the left m bandlet (circle at the end of the white dashed line) is mispositioned. The open asterisk indicates the ectodermal bandlets on the left side. The yellow dashed line outlines the well-developed right germinal band. The M_L_ teloblast was co-injected with the rhodamine dextran tracer and *Hau*-*netrin-1* mRNA immediately after its birth in early stage 5; the image was taken in early stage 8. (B) The m bandlet (red) had a normal morphology following control injection. (C) The m bandlet also exhibits a normal morphology after the M_L_ was co-injected with tracer and *Hau*-*netrin-*2 mRNA in early stage 5. (D) Quantification of the experiments shown in panels A-C. (E) The m bandlet (red) is detached from the ectodermal bandlets (green) in a mid-stage 8 embryo, whose M_L_ was co-injected with the rhodamine dextran tracer and *Hau-netrin-1* mRNA and NOPQ_L_ with a fluorescein dextran tracer in early stage 5. (F) In the control mid-stage-8 embryo, the m bandlet (red) aligned with the four ectodermal bandlets (green) on its top. (G) The right m bandlet (green) converged toward the left m bandlet, which misexpressed Hau-netrin-1 (red) in the prospective dorsal territory. The anterior tips of both bandlets were mispositioned. M_L_ teloblast was co-injected with *Hau-netrin-1* mRNA and rhodamine dextran in and M_R_ with fluorescein dextran in stage 5. (H) The left and right m bandlets exhibited normal bilateral configuration in stage 8, following control injections in stage 5. (I) Quantification of the experiments shown in panels G and H. The non-cell autonomous effect of Hau-netrin-1 misexpression in the left m bandlet was scored by the mispositioning of the right m bandlet, which did not misexpress Hau-netrin-1, in 39% of the operated embryo. (J and J’) *Hau-netrin-1* mRNA was co-injected with rhodamine dextran after the M_L_ teloblast had produced several blast cells in stage 6b. The left m bandlet (red) succeeded in entering the germinal band in the resulting stage 8 embryo, but the germinal band bends dorsally in the section where ectopic Hau-netrin-1 is produced (white arrowhead). The right germinal band also veered dorsally (yellow arrowhead), though not as severely as the left germinal band. Panel J is from dorsal view; J’ is from the lateral view, with dorsal to the right. (K and K’) Normal position of the left m bandlet (red) in a mid-stage-8 embryo treated with control injection. Panel K is from dorsal view; K’ is from the lateral view, with dorsal to the right. (L) Quantification of the experiments shown in panels J, J’, K and K’. (M) The left m bandlet (red; labeled by tracer injection into the M_L_ teloblast) entered the germinal band, and the germinal band bends dorsally (white arrowhead) in an embryo whose left o, p and q bandlets misexpress Hau-netrin-1 (green) by co-injecting mRNA and tracer into the OPQ_L_ proteloblast. The unoperated right germinal band was also dorsally positioned (yellow arrowhead). (N) The germinal band morphology was normal in an embryo treated with control injections. (O) Quantification of the experiments shown in panels M and N. Scale bar: 150 μm.

Since netrin is a secreted signaling ligand, we next asked whether Hau-netrin-1 misexpression in the left m bandlet can affect the migration of the right m bandlet. To address this question, we co-injected *Hau-netrin-1* mRNA with a tracer into the left M teloblast and a different tracer into the right M teloblasts. In 39% of the operated embryos, the left and right germinal bands converged toward the dorsal midline, instead of the ventral midline, in early stage 8 (Figures 5G and 5I). In contrast, all control embryos injected with tracers alone were normal in the same stage (Figures 5H and 5I). Further time-course analysis showed that the anterior ends of the left and right m bandlet failed to separate during stage 7 in embryos mis-expressing Hau-netrin-1 in the left m bandlet (Figure 6), indicating that Hau-netrin-1 acts as an attractive signal for the m bandlets, which expresses Hau-DCC2, during germinal band assembly. This is consistent with that the expression of the DCC-type receptor encodes a positive chemotaxis behavior toward the netrin gradient in the responding cells (Chan et al., 1996; de la Torre et al., 1997; Hong et al., 1999).

**Figure 6.**
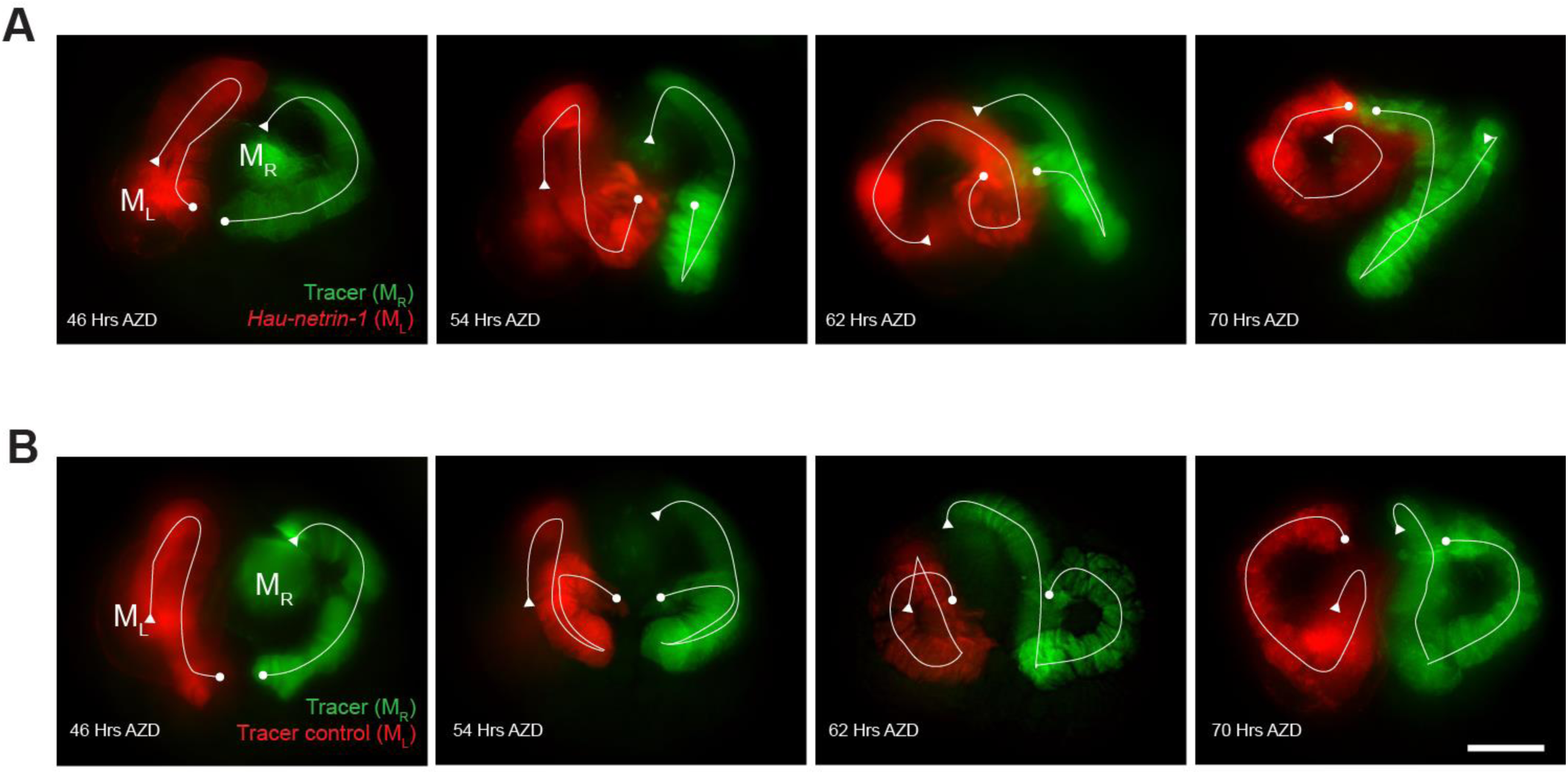
Time series of mesoderm morphogenesis in embryos ectopically expressing Hau-netrin-1 in the left m bandlet (A) and normal embryos (B). (A) In embryo misexpressing Hau-netrin-1 in the left m bandlet, the anterior end of the left and right m bandlets remain in close physical association throughout the course of observation. The m bandlet morphology at 46 hours AZD was similar to the control at the same time point but became increasingly aberrant in later developmental stages. These embryos were co-injected with rhodamine dextran tracer and *Hau-netrin-1* mRNA in the M_L_ teloblast and then injected with fluorescein dextran in the M_R_ teloblast. (B) The normal configurations of the m bandlets at 46 hours AZD, 54 hours AZD, 62 hours AZD, and 70 hours AZD. These control embryos were injected with rhodamine dextran tracer in the M_L_ teloblast and fluorescein dextran tracer in the M_R_ teloblast. Each of the m bandlets is labeled with a white line; the dot at one end of the line marks the anterior tip of the bandlet; the triangle at the other end marks the posterior extreme of the bandlet, where the blast cells arise from the teloblast. Scale bars: 100 μm.

### Hau-netrin-1 is involved in two distinct tissue migration events during leech gastrulation

Since the expression of Hau-netrin-1 in the n bandlet persisted throughout the entire course of gastrulation, we next went on to determine if Hau-netrin-1 has any additional role in later stages of gastrulation. To avoid disrupting germinal band assembly, we injected *Hau*-*netrin-1* mRNA into the M teloblast in stage 6b. In these embryos, the ectopic Hau-netrin-1 was limited to the more posterior part of the m bandlet, and the germinal band assembly would have been completed before Hau-netrin-1 misexpression reached an appreciable level (Gline et al., 2011). As expected, the entrance of m bandlet into the germinal band was normal in all operated embryos. However, the epibolic migration of the germinal band was disrupted, leading to an “exogastrula” phenotype in which the left and right germinal bands remained in the dorsal territory of the embryo in 30% of the observed cases (Figures 5J, 5J’ and 5L). In these abnormal embryos, the ectodermal bandlets and the m bandlet were bonded together to form a coherent germinal band but were misplaced, indicating that they migrate as a unit. No such phenotype was observed in the control embryos injected with tracer only (Figures 5K, 5K’ and 5L). Together, our results suggested that Hau-netrin-1 has two distinct roles in mesoderm migration during gastrulation. First, it guides the entrance of the m bandlet into the germinal band in the initial phase of gastrulation. Second, it drives the epiboly of the germinal band in the later stage of gastrulation.

### Ectoderm-derived netrin gradient drives germinal band epiboly

Given that *Hau-netrin-1* was expressed in the n bandlets, the ventralmost ectodermal bandlet, a ventral-to-dorsal gradient of Hau-netrin-1 should arise in the germinal band to guide its ventralward migration during epiboly. To test this hypothesis, the ventral-to-dorsal gradient of Hau-netrin-1 was disrupted by ectopically expressing Hau-netrin-1 in the o, p, and q bandlets. In 31% of the operated embryos in which the OPQ”_L_ proteloblast was injected with *Hau-netrin-1* mRNA, the labeled left m bandlet, together with the overlying ectodermal bandlets, veered toward dorsal midlines, and the right germinal bands also appeared affected and moved toward the midlines (Figures 5M and 5O). The organization and positioning of the germinal band were normal in all control embryos (Figures 5N and 5O). Thus, we concluded that the netrin gradient on the ectodermal layer of the germinal band drives its epibolic movement toward the prospective ventral midline.

## Discussion

Here, we studied the expression and function of the evolutionarily conserved midline marker netrin during gastrulation of the leech *Helobdella*. The late-stage expression patterns of the two leech netrin homologs generally agree with the conventional interpretation of netrin’s roles as a chemotactic factor for axon guidance and organogenesis. Expression of *Hau-netrin*-*2* at the mesodermal midline may represent its canonical function as a midline landmark, and its expression in the nephridia suggested involvement in the morphogenesis of nephridial tubule. The ganglionic expression of *Hau-netrin-1* is indicative of possible roles in axon guidance and the morphogenesis of segmental ganglia. However, the expression of *Hau*-*netrin-1* at the onset of gastrulation is highly unusual and of great interest, given that netrin has never been implicated in gastrulation of any metazoan species. The netrin-expressing cells in leech gastrula would eventually give rise to the midline structure, but netrin expression begins before these cells assume such a position, suggesting a heterochronic shift of netrin expression, relative to that observed in the ventral midline of the polychaete larva (Denes et al., 2007; Lauri et al., 2014), has occurred in the evolutionary lineage leading to the leech. This heterochronic shift is best interpreted as resulting from the evolution of direct development by accelerating trunk tissue development in the clitellate ancestor (Kuo, 2017).

Furthermore, accelerated trunk development also brought about a unique form of gastrulation in clitellate annelids, in which germinal bands – the primordial trunk tissue – undergo epibolic migration and posterior growth simultaneously. Our results suggested that netrin is involved in the formation of the germinal band. Specifically, we demonstrated (1) that Hau-netrin-1 is expressed by the n bandlet before germinal band assembly, (2) that the N lineage is required for the entrance of the m bandlet, which expresses Hau-DCC2, into the germinal band, and (3) that Hau-netrin-1 can guide the migration of the m bandlet. Together, these suggested that Hau-netrin-1 expressed by the n bandlet may attract the m bandlet toward the superficial ectodermal bandlets to assemble the germinal band. However, given the relatively low penetrance in both ablation and misexpression experiments and the lack of observable abnormal phenotype in the attempted knockdown of Hau-netrin-1 and the over-expression of dominant-negative Hau-DCC2 (data not shown), we anticipated that additional factors would contribute to germinal band assembly in parallel to Hau-netrin-1. The mechanical stress generated by the expansion of growing tissue and the space constraint imposed by the very narrow blastocoel of the *Helobdella* embryo can push the m bandlet toward the ectodermal bandlets; other chemotactic signals and cell sorting by differential cell adhesion may also be involved in the movement of the m bandlet into the germinal band.

After the germinal band has formed, Hau-netrin-1, which continues to be expressed by cells of the n bandlet, forms a ventral-to-dorsal gradient across the germinal band. When this gradient was disrupted by Hau-netrin-1 misexpression in the lateral and dorsal ectoderm, the germinal band failed to migrate ventrally and remained in the dorsal region, resulting into an exogastrula phenotype. This suggested that Hau-netrin-1 may further guide epibolic migration of the germinal band after the germinal band had been assembled. Since the m bandlet serves as the scaffold for the attachment of ectodermal bandlets and is central to the organization of the germinal band (Blair, 1982; Zackson, 1984), it is likely that Hau-netrin-1 guides germinal band migration by attracting the m bandlet, which in turn carries the ectodermal bandlets with it; this resembles the carrot-on-a-stick trick. Nonetheless, we cannot completely rule out the possibility that Hau-netrin-1 also actively attracts other ectodermal bandlets, since *Hau*-*DCClike* is expressed by the ectodermal bandlets in the germinal band. In any case, a more complex scenario is expected for the mechanism of leech gastrulation, provided that mechanical forces created by the expanding micromere cap and the elongating germinal band also contribute to this process (Keller, 2006; Smith et al., 1996). It is also noteworthy that, in the more advanced stages of germinal band development, the leading edge of the mesoderm moves ahead of the ectoderm, violating the netrin gradient. Thus, we speculated that the carrot-on-a-stick mechanism might have a more critical role in the initial phase of epiboly than in the later stages. Taken together, changing combinations of chemotactic signals and oriented cell divisions may drive epibolic migration of the germinal bands in different stages of gastrulation.

In this study, we have revealed the unusually early expression of the leech netrin genes in the midline precursor cells and showed that netrin signal could be employed to guide cell migration during gastrulation. These novel functions of netrin are likely evolutionarily derived, specifically in the clitellates – the clade that includes most of the annelid species found in freshwater and terrestrial habitats, such as earthworms and leeches. Despite the unavailability of data outside of the leeches, netrin signaling may have a similar role in the gastrulation of other clitellates, since the same form of early embryogenesis and gastrulation as described in *Helobdella* is shared among the clitellates and likely a clitellate apomorphy (Anderson, 1973).

What was the selective advantage of recruiting netrin to guide mesoderm migration in clitellate gastrulation? It is reasonable to assume that some form of cell-cell communication is required to coordinate such a complex and reproducible pattern of cell migration during gastrulation. Indeed, the axial patterning morphogens, such as BMP, Nodal, and FGF, have been shown to regulated zebrafish epiboly (Heisenberg and Solnica-Krezel, 2008). Although many such signals were shown to work by modulating cell motility rather than acting as a chemotactic factor, there are a few cases where chemotactic effects of selected signaling molecules have been documented in gastrulation (Adomako-Ankomah and Ettensohn, 2013; Yang et al., 2002). Above all, none of these growth factors is purely chemotactic as they all have dual roles as a morphogen and a chemotactant. Together, in those embryo patterned by morphogen, the morphogen systems themselves are responsible for orchestrating cell movements in different regions of the gastrulating embryo.

In contrast, there is no global morphogen in an embryo with a fixed cell lineage, *e.g.,* that of the nematodes, the ascidians, and the clitellates. However, unlike the leech embryo, no chemotactic factor is expressed during gastrulation in the simpler and smaller nematode embryos. This can be readily explained by the absence of long-distance cell migration in early nematode embryos (Montell, 1999). Therefore, we postulate that the co-option of chemotactic factors to guide cell migration in gastrulation may represent a unique strategy for maintaining the precise and robust long-distance cell movements in gastrulation of the larger and more complex clitellate embryos. It is interesting to note that the clitellate mode of embryonic development first emerged in part as a consequence of increasing egg size in the clitellate ancestor (Anderson, 1973; Kuo, 2017). Thus, the accelerated expression of midline netrin may mechanistically connect the evolution of cell lineage-dependent embryogenesis, the evolution of direct development, and the evolution of epibolic gastrulation in the clitellates. This study has highlighted the potential of the clitellate embryos as a model system for identifying and characterizing the molecular changes underlying the evolution of axial patterning and gastrulation in animal development.

## Materials and Methods

### Animal

Embryos of *Helobdella austinensis* were used in this study (Kutschera et al., 2013). The adult leeches were cultured in 1% artificial seawater at room temperature and were fed with chironomid larvae five to six times a week. Embryos were retrieved from cocoons attached to the ventral surface of their mother and were cultured at 25°C in the modified HL medium (17.3 mM NaCl, 1.2 mM KCl, 2 mM MgCl_2_, 8 mM CaCl_2_, 1 mM maleic acid, pH 6.6) supplemented with antibiotics (10 μg/mL tetracycline; 50 U/mL penicillin-streptomycin).

### Gene identification and molecular cloning

*In silico* gene identification was carried out by BLAST searches of the whole genome assembly of *Helobdella robusta* (Simakov et al., 2013), a sibling species of *H. austinensis*. Two netrin ligands, three DCC-type receptors, and two Unc5-type receptors were identified in the *H. robusta* genome (Table S1). Using the *H. robusta* sequences, PCR primers (Table S2) were designed to amplify cDNA fragments of the complete set of netrin ligands and receptors in *H. austinensis* (*Hau-netrin-2*, *Hau-netrin-1*, *Hau-DCC1*, *Hau-DCC 2*, *Hau-DCC-like*, *Hau-Unc5A,* and *Hau-Unc5B*). A pool of mixed-stage *H. austinensis* cDNA was used as the template, and PCR reaction was performed using the proof-reading Phusion DNA polymerase (ThermoFisher). 3’-termini A overhangs were introduced to the amplicons by a brief treatment with non-proofreading Taq DNA polymerase in the presence of dATP. The treated amplicons were gel extracted and inserted into pGEM-T (Promega) or pCRII (ThermoFisher). The cloned cDNA fragments were sequenced to ensure their molecular identities. These plasmids were used as templates for riboprobe synthesis in the subsequent *in situ* hybridization experiments. If necessary, the cDNA sequences were further extended by the rapid amplification of cDNA end (RACE) method by using FirstChoice RLM-RACE Kit (ThermoFisher). Gene-specific primers used in the RACE PCR reactions are listed in Table S3. The full-length coding sequence were assembled from the cloned cDNA fragments, and the full-length coding cDNA of *Hau-netrin-1*, *Hau-netrin-2* were PCR amplified from *H. austinensis* embryonic cDNA using Phusion DNA polymerase (ThermoFisher). The primer pairs used in these PCR reactions are listed in Table S4. The cDNA fragments containing the full-length coding region was then subcloned into pCS107 vector. The resulting plasmids were used as templates for *in vitro* synthesis of mRNA.

### Phylogenetic analysis

To determine gene orthology for the leech netrin ligands and receptors, phylogenetic analyses were performed for each of the three classes of genes, namely netrin, DCC and Unc5. Sequences of the leech cDNA clones were conceptually translated to amino acid sequences. Amino acid sequences of netrin, DCC and Unc5 homologs from other metazoan species were retrieved from the GenBank database. Alignments of netrin, DCC, and Unc5 families were generated using MUSCLE algorithm (Edgar, 2004) implemented in the MEGA software (Kumar et al., 2016). Specific conserved domains in the alignments were further identified by NCBI Conserved Domain Search (Marchler-Bauer et al., 2015). Alignments that contain all conserved domains were obtained by removing residues outside of the conserved domains. For netrin, an alignment that contains only the laminin_N domain was produced for the netrin alignment. Phylogenetic trees were constructed either by the Neighbor-Joining method, using MEGA software (Kumar et al., 2016), or by the Maximum-Likelihood method, using RAxML software (Stamatakis, 2014). 500X bootstrap was performed for each tree. Further details regarding each specific phylogenetic tree are given in the legends of Figure S1.

### Gene expression analysis

Developmental expression of the leech netrin and netrin receptor genes was determined by whole-mount *in situ* hybridization (WMISH). To perform WMISH, embryos of the specific developmental stage were fixed in a 1:1 mixture of 8% formaldehyde and PBS at 4°C overnight and stored in methanol. Standard single-color chromogenic WMISH was carried out as previously described (Weisblat and Kuo, 2009). The color reaction was carried out using BM Purple phosphatase substrate (Roche). Following the color reaction, the specimens were dehydrated and stored in ethanol. The specimens were cleared and mounted in 70% glycerol for imaging. The images were taken using an Olympus DP80 CCD camera mounted on an Olympus BX63 microscope. Digoxigenin-labeled riboprobes were *in vitro* transcribed from DNA templates produced by PCR amplification of cDNA fragment from the plasmids containing the specific cDNA fragments.

The protocol for fluorescence WMISH (FWMISH) is similar to standard WMISH. In FWMISH, PreHyb solution was supplemented with 5% dextran sulfate (Sigma), and a higher probe concentration than the standard WMISH was used. The optimal probe concentration was empirically determined for each probe and each developmental stage. In double FWMISH, the two riboprobes were labeled with digoxigenin and DNP respectively. Following hybridization, peroxidase-conjugated anti-digoxigenin antibody (Roche), applied at 1:1000 dilution in blocking solution, was used to tag the digoxigenin-labeled probe. Tyramide signal amplification (TSA) method was used to tag fluorophore to the location of probe binding. Following washes in the borate buffer (100 mM borate, pH 8.5; 0.1% Tween-20), specimens were placed in freshly made TSA reaction solution (0.5 mg/mL fluorophore-conjugated tyramide, 2% dextran sulfate, 500 μg/mL 4-iodophenol, and 0.003% H_2_O_2_, in 0.1 M borate). The conjugated tyramide was home-synthesized from fluorophore-conjugated reactive amine (ThermoFisher) and tyramide (Sigma) following a previously described protocol (Hopman et al., 1998). Following TSA reaction, the specimens were washed extensively to remove the background resulting from the unreacted fluorophore molecules. When double FWMISH was performed, the specimens were treated briefly in acidic glycine buffer (0.1 M glycine, pH2.0; 0.1% Tween-20) to remove the peroxidase-conjugated anti-digoxigenin antibody from the specimens. The specimens were then labeled with a peroxidase-conjugated anti-DNP antibody (Perkin Elmer). A different fluorophore was used to carry out the second TSA reaction. The specimens embryos were cleared and mounted in 80% buffered glycerol containing 4% n-propyl gallate for imaging.

### mRNA synthesis

*In vitro* synthesis of capped RNA was performed by using mMESSAGE mMACHINE SP6 kit (ThermoFisher). DNA template for mRNA synthesis was prepared by linearizing the pCS107-derived plasmid by AscI restriction digestion. The synthesized mRNA was further polyadenylated *in vitro* by using Poly(A) Tailing Kit (ThermoFisher) to improve the efficiency of misexpression in early developmental stages. The synthesized mRNA were purified and concentrated by using RNA Clean & Concentrator-5 column (Zymo Research), aliquoted and stored in −80°C.

### Embryological experiments

Lineage tracing experiments were performed by microinjection of dextran-conjugated tetramethylrhodamine (RD; ThermoFisher “Fluoro-Ruby”) or fluorescein (FD; ThermoFisher “Fluoro-Emerald) into an identified blastomere. RD or FD was injected at 15 mg/mL in pipette, and 0.1% Phenol Red (Sigma) or 0.25% FastGreen (Sigma) was used as a monitoring dye for microinjection. For cell ablation experiments, 5 mg/mL DNase I (Roche) was co-injected with a fluorescent dextran tracer into an identified cell. Injected DNaseI prevents further division of the injected cell by quenching the cytoplasmic pool of G-actin. In misexpression experiments, *in vitro* synthesized mRNA was co-injected with fluorescent dextran tracer. The mRNA was injected at 800 ng/mL in pipette. The amount of injectant delivered was estimated to be 1/500 of the cell volume. Microinjection was performed as previously described (Weisblat and Kuo, 2009).

The injected embryos were raised to the specific stage and fixed for documentation. Fixed specimens were counterstained with DAPI (Sigma) and mounted in buffered glycerol containing 4% n-propyl gallate for fluorescence imaging. Fluorescence micrographs were generated by digital deconvolution of wide-field fluorescence images captured with an Olympus DP-80 digital camera mounted on an Olympus BX-63 fluorescence microscope, using cellSens software (Olympus) or by confocal microscopy using a Leica SP5 system.

Each set of embryological experiments triplicated, and data were pooled for presentation. Control experiments were performed in parallel to the designated ablation or misexpression experiments using embryos of the same cohort. Chi-square test was employed for testing the statistical significance between the treated group and the corresponding control.

## Acknowledgments

We thank the staffs of TechComm, College of Life Science, NTU for assistance in confocal imaging. This work was supported by Ministry of Science and Technology, Taiwan (103-2311-B-002-019 and 104-2311-B-002-025-MY2 to DHK).

---

**Figure S1.**
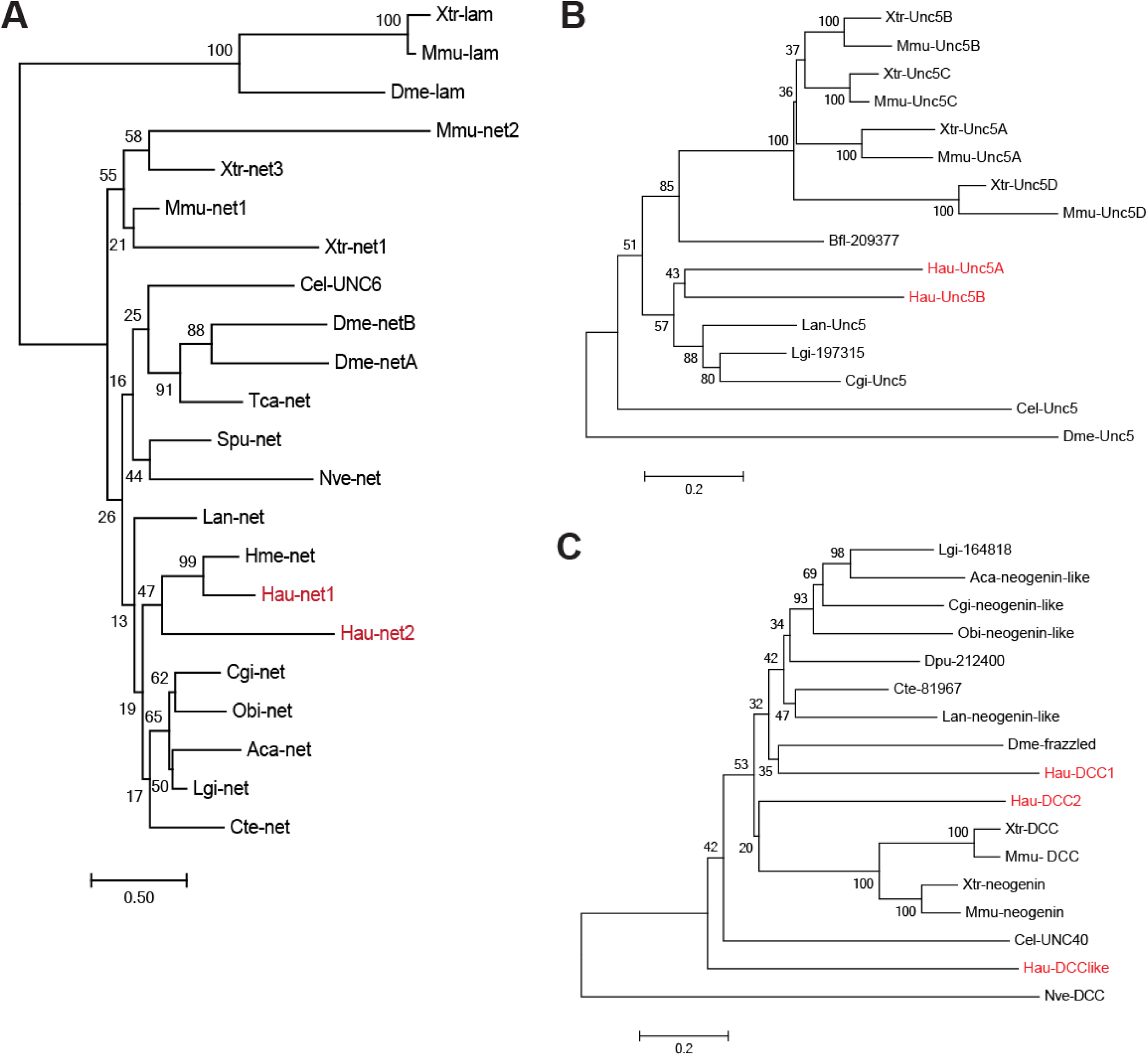
Phylogenetic trees of the netrin family, DCC family, and Unc5 family. A three-letter prefix denotes the species from which the sequence originates from. Aca: *Aplysia californica*; Bfl: *Branchiostoma floridae*; Cel: *Caenorhabditis elegans*; Cgi: *Crassostrea gigas*; Cte: *Capitella telata*; Dme: *Drosophila melanogaster*; Dpu: *Daphnia pulex*; Hau: *Helobdella austinensis*; Hme: *Hirudo medicinalis*; Lan: *Lingula anatine*; Lgi: *Lottia gigantea*; Mmu: *Mus musculus*; Nve: *Nematostella vectensis*; Obi: *octopus bimaculoides*; Tca: *Tribolium castaneum*; Xtro: *Xenopus tropicalis*. 1,000 X bootstraps were performed for each tree, and the bootstrap support value is labeled on each node. (A) A LG+G Maximum-Likelihood tree of the netrin genes generated from an amino acid alignment of the Laminin_N domain from selected metazoan netrin homologs and the laminin outgroups. Accession numbers of the sequences used in this alignment, other than those from *Helobdella austinensis*, are listed below. Aca-net: AJD80259.1; Cel-UNC6: NP_509165.1; Cte-net: ELU09550.1; Dme-netA: AAB17548.1; Dme-netB: AAB17547.1; Hme-net: AAC83376.1; Lan-net: XP_013408519.1; Lgi-net: 201560, JGI; Mmu-net1: AAC52971.1; Mmu-net3: AAH94362.1; Nve-net: ABF61780.1; Obi-net: XP_014780148.1; Tca-net: XP_008199140.1; Xtr-net: AAI21569.1; Xtr-net3: XP_002932466.2; Dme-lam: NP_476617.1; Mmu-lam: EDL38338.1; Xtr-lam: XP_004915294.1. (B) A Neighbor-Joining tree of Unc5 genes generated from an amino acid alignment of all conserved domains of selected metazoan homologs. Bfl-209377: 209377, JGI; Cel-UNC5: NP_500823.1; Cgi-UNC5: EKC29674.1; Dme-UNC5: AAF58143.2; Lan-UNC5: XP_013412208.1; Lgi-197315: 197315, JGI; Mmu-UNC5A: AAH58084.1; Mmu-UNC5B: AAH57560.1; Mmu-UNC5C: AAI15773.1; Mmu-UNC5D: AAI44785.1; Xtr-UNC5A: NP_001093674.2; Xtr-UNC5B: XP_002939005.2; Xtr-UNC5C: XP_004911160.1; Xtr-UNC5D: XP_002933002.2. (C) A Neighbor-Joining tree of the DCC genes generated from an amino acid alignment of the fibronectin domains of selected metazoan homologs. Cel-UNC40: NP_491664.1; Cgi-neogenin-like: XP_011437571.1; Cte-17814: 17814, JGI; Dme-frazzled: AAC47314.1; Dpu-212400: 212400, JGI; Lan-neogenin-like: XP_013387952.1; Lgi-164818: (164818, JGI; Mmu-DCC: NP_031857.2; Mmus-neogenin: NP_032710.2; Nve-DCC: AIZ68366.1; Obi-neogenin-like: XP_014777890.1; Xtr-DCC: XP_004910423.1; Xtr-neogenin: XP_012821603.1.

**Figure S2.**
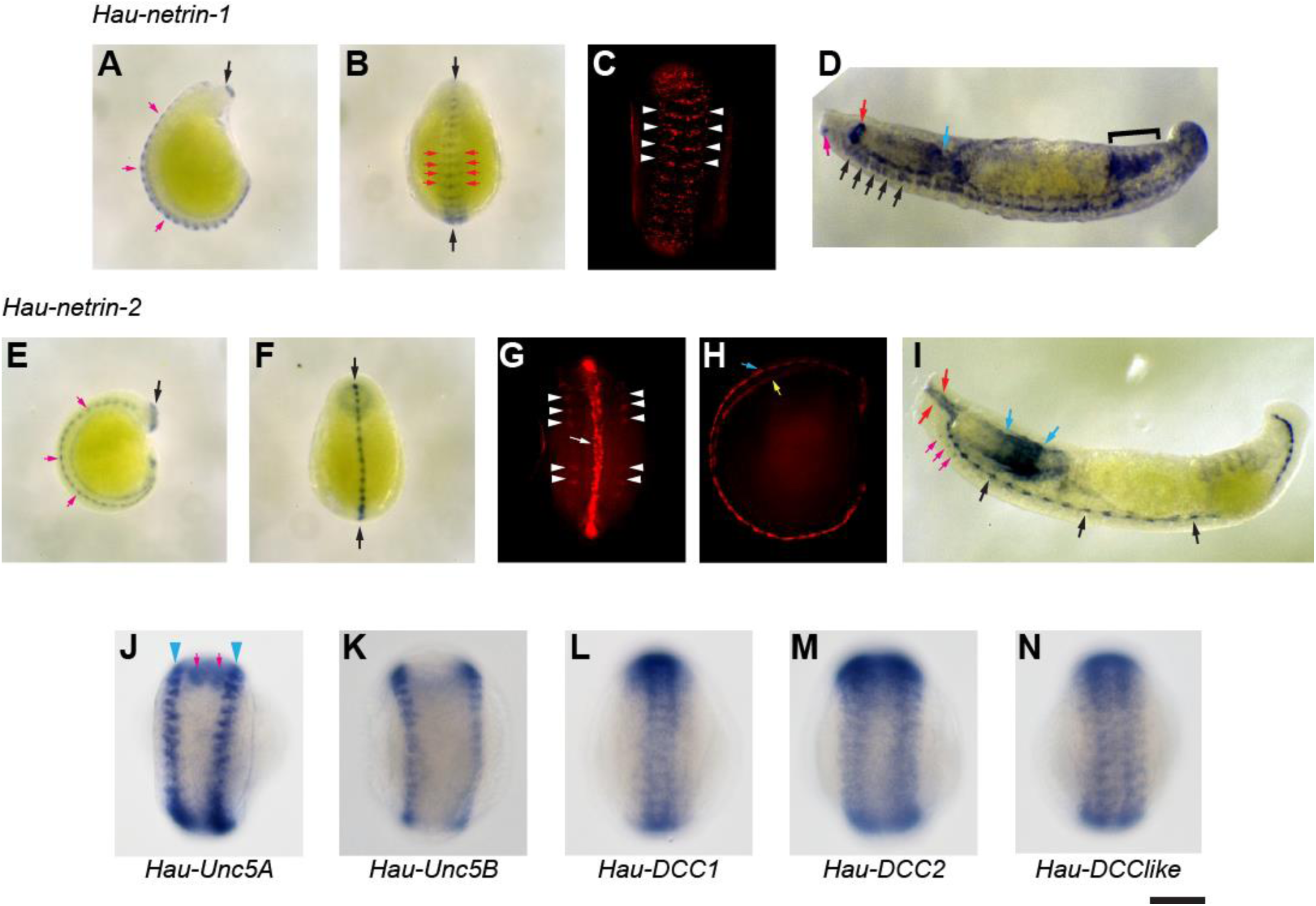
Expression patterns of netrins and receptors in late embryonic development of the leech revealed by WMISH. (A) Lateral view of a stage-9 embryo stained for *Hau-netrin-1*. Anterior is to the top; dorsal to the right. Black arrow indicates a ring of *Hau-netrin-1* expressing tissue near the anterior end of the developing proboscis. Pink arrows point to ventral ectodermal *Hau-netrin-1* expression in the trunk. (B) Ventral view of the same embryo shown in panel A. In addition to the discontinuous expression along the ventral midline (arrow), *Hau-netrin-1* expression also extends bilaterally in each segmental ganglion (red arrows). (C) Fluorescent *Hau-netrin-1* WMISH showed that the *Hau-netrin-1* domain extends bilaterally further (arrowheads) while the midline expression becomes less prominent in early stage 10. Ventral view. (D) *Hau-netrin-1* has a complex expression pattern in early stage 11. It is expressed in the segmental ganglia (black arrows indicate the five anteriormost ganglia), a ring in the proboscis sheath (red arrow), the base of proboscis (blue arrow), the intestine (bracket), and the anterior part of the front sucker primordium (pink arrow). Lateral view; anterior is to the left; dorsal to the top. (E) WMISH of *Hau-netrin-2* revealed its expression in the mesodermal midline (pink arrow) and a ring near the base of proboscis (black arrow) in early stage 9. Lateral view; anterior is to the top; dorsal to the right. (F) Ventral view of the same embryo shown in panel E. Arrows indicate mesodermal midline expressing *Hau-netrin-2*. (G) Fluorescent *Hau-netrin-2* WMISH of an early stage-10 embryo (ventral view). In addition to the mesodermal midline (arrow), *Hau-netrin-2* is also expressed in the developing nephridia (arrowheads). (H) Lateral view of the same embryo in panel G showed *Hau-netrin-2* expression in the midline of both the body-wall mesoderm (blue arrow) and visceral mesoderm (yellow arrow). (I) *Hau-netrin-2* is expressed in the midline (arrows), selected cells in the rostral segmental ganglia (pink arrows), the wall of proboscis chamber (red arrows), the posterior section of the proboscis and proboscis sheath (blue arrows) in stage 11. (J) *Hau-Unc5A* is expressed in the dorsal region of the germinal plate (blue arrowheads) of the stage-9 embryo; this is likely the continuation of dorsal germinal plate expression in stage 8 (Figure 2M). (K) Similar to stage 8, *Hau-Unc5B* expression remains in the dorsal margin of the germinal plate in stage 9. (L-N) The expression patterns of the DCC-family receptors (L: *Hau-DCC1*; M: *Hau*-*DCC2*; N: *Hau-DCClike*) in stage 9 are similar to their respective patterns in stage 8 (*Hau*-*DCC1*, Figure 2Q; *Hau-DCC2*, Figure 2S; *Hau-DCClike*, Figure 2U). Scale bar: 150 μm in panels C, G, H, and J-N; 200 μm in panels A, B, E, and F; 75 μm in panels D and I.

**Table S1.**
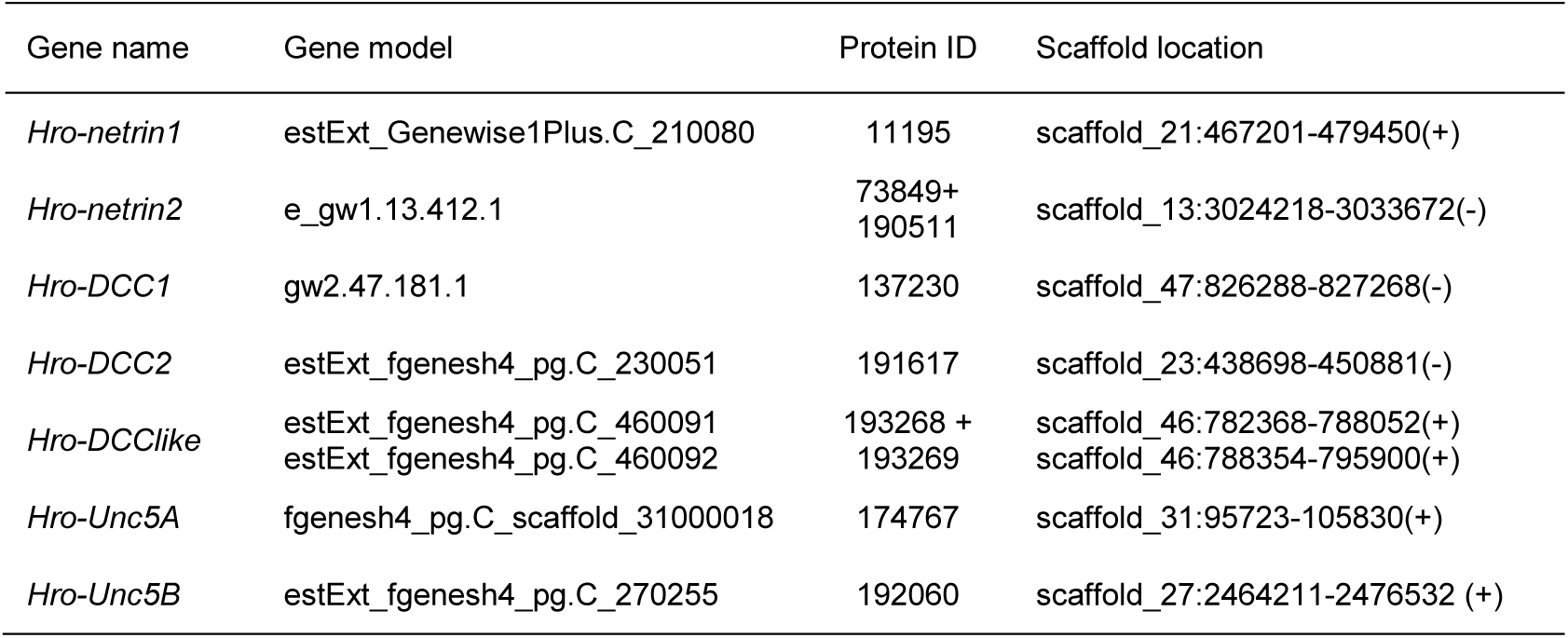
Gene IDs of the leech netrins and netrin receptors in the JGI database

**Table S2.**
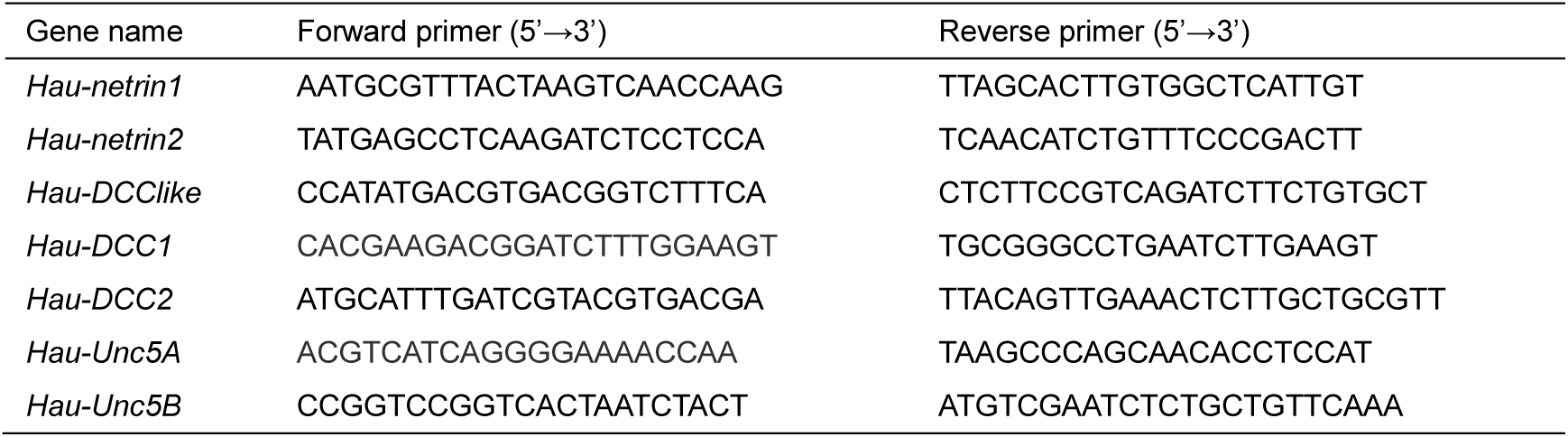
Primers pairs for cDNA cloning

**Table S3.**
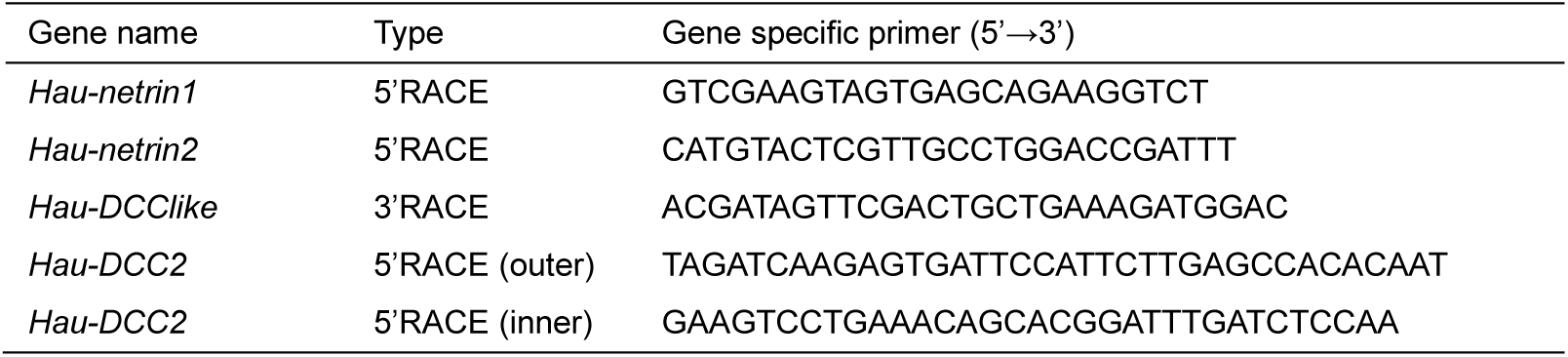
Gene-specific primers for RACE

**Table S4.**
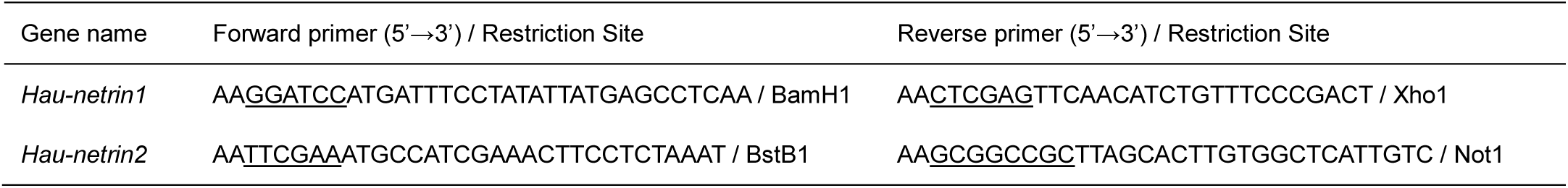
Primer pairs for overexpression construct of *Hau-netrin1*and *Hau-netrin2*

